# A study of conventional histological parameters in excision specimens of breast cancer over a 7 year period and association with immunohistochemical categories

**DOI:** 10.1101/2020.06.15.149815

**Authors:** Parikshit Sanyal, Anshuman Singh, Prosenjit Ganguli, Sanghita Barui

## Abstract

**Background:** Along with conventional histological examination, immunohistochemistry (IHC) is a useful adjunct to assessment of a breast cancer excision specimen. Previous studies have shown differences in behavior of neoplasms depending on their histopathological as well as immunohistochemical categories; in particular, triple negative breast cancers (on IHC) show the worst prognosis.

**Objectives:** To find association, if any, within conventional histopathological characteristics (size, grade, stage, mitotic count, desmoplasia, dense inflammatory infiltrate, lymphovascular invasion) and between the conventional parameters and immunohistochemical categories of breast cancer, in both primary and post neo adjuvant chemotherapy (NACT) specimens.

**Methods:** 177 breast cancer excision specimens examined over last 7 years were assessed retrospectively, their histopathological parameters were recorded. In cases where immunohistochemistry was performed (N=108) the specimen was placed in one of the immunohistochemical categories: Luminal A, Luminal B, Her2 and Triple negative cancers. The data was then analysed by standard statistical methods.

**Results:** No statistically significant association was found between the histopathological parameters and IHC category was. However, a strong correlation was seen between lymphovascular invasion within the primary tumor and increasing lymph node involvement. There was also a reduction in ER and PR expression in neoplasms post NACT, while HER2 expression remained largely unchanged.

**Conclusion:** There might be additional genetic subtypes underlying the immunohistochemical phenotypes which determine the morphological characteristics of the neoplasm.

## Introduction

Breast cancer is the commonest cancer of Indian women [1], creeping ahead of cervical cancer in 2012. The pattern of breast cancer in India differs from the West in a few significant aspects, one of them being that triple negative breast cancers, i.e. not expressing Estrogen Receptors (ER), Progesterone Receptors (PR) and Human Epidermal Growth Factor Receptor 2 (HER2), are much more commoner in India than the west [2]. This is a critical difference, because cancers which express none of these markers (‘triple negative’) have been proven to have the worst prognosis, followed by the ER/ PR negative and Her2 positive subtype [3]. Because of availability of a monoclonal antibody against HER2, trastuzumab, expression of HER2 has also become a predictive marker in breast cancer. With the widespread availability of immunohistochemistry, all breast cancers are now being routinely tested for expression of the two hormone receptors ER and PR, and the growth factor receptor HER2.

Conventionally, the pathologist examines the breast cancer specimen within the ambit of a few gross morphological and microscopic parameters. This include the tumor size on gross, which is the primary determinant of the ‘T’ stage of the cancer. The histological grade is determined from the Scarff Bloom Richardson (SRB) score [4], which takes into account the amount of tubule formation within the tumor, the subjective assay of nuclear pleomorphism and the proportion of dividing cells in the tumor. The ‘N’ stage of a tumor is determined by the number of lymph nodes involved by the cancer. Lymph node staging can only be done when axillary dissection has been carried out concurrently with a Modified Radical Mastectomy (MRM. Within the primary neoplasm, the subjective assessment of presence or absence of a dense inflammatory infiltrate around the neoplasm, and that of desmoplastic (fibrotic) reaction around the neoplasm have also been reported as useful adjuncts to predict neoplasm behavior. The College of American Pathologists recommends recording of skin involvement by the tumor, if any, both on gross and microscopy [5]

It is well known that Hormone Receptors and HER2 are of prognostic significance in breast cancer. The present study aims to find correlation, if any, between conventional pathological parameters of a cancer and immunohistochemical categories, which might shed light on the pathogenesis of the different cancer subtypes based on marker expression, as well as assist the pathologist and the clinician in predicting neoplasm behavior when immunohistochemical facilities are not available.

## Method

A retrospective, analytical study design was chosen. Initial ethical clearance for accessing Hospital records was obtained from the Institutional Ethical Committee. The histopathologic records maintained in the Laboratory Information System (LIS) over last 7 years was mined and analysed. Between 2010 to 2017, 183 core biopsies of breast, 131 Modified Radical Mastectomies (MRM), 25 excisions less than total mastectomy or ‘lumpectomy’ (LM), 17 Wide Local Excisions (14.04%) and 4 Simple Mastectomies (SM) were examined in this Department. 01 (one) set of slides and blocks was received from another hospital for consultation, which was also included in the study. A total of 177 cases of excisions and mastectomies were analysed; 34 of these cases were resected post neo adjuvant chemo therapy (NACT). 183 (one hundred eighty three) core biopsies from breast was also performed in this period, and 126 (68.85%) of these biopsies turned out to be malignant. 19 of these patients who continued treatment in this Hospital were traced from the archives and their ER, PR status post NACT and mastectomy was recorded.

### Inclusion criteria

All primary neoplasms of the breast in female patients in the years 2011-2017; including

1. Those excision specimens which were received post NACT
2. Those biopsy specimens which were reported malignant, treated with NACT, and underwent some form of excision

### Exclusion criteria

Metastatic carcinomas in breast reported in this period were excluded from this study.

### Data collection and analysis

After the initial survey of LIS, an electronic proforma was prepared for data collection, which included the patients’ demographic characteristics, type of specimen received, subtype of cancer identified, morphologic information (size, skin involvement), pathologic stage (both T and N stage) histologic features (dense infammatory infiltrate, desmoplasia, lymphovascular invasion), and whether the specimen was primary neoplasm or post NACT.

The collected data was tabulated in a database, and the Sequential Query Language (SQL) was used to mine this data to find out correlation between the variables. Standard statistical methods were used for analysis. Statistical tools like Libreoffice Calc, Python with Numpy and Scipy library were used for analysis.

The neoplasms were divided in the following immunohstochemical subtypes for the purpose of analysis, as per the criteria proposed in Carolina Breast Cancer study [6] [7] (Table 1).

**Table 1.** Division into immunohistochemical subtypes

|  |  |
| --- | --- |
| Luminal A | ER and/or PR positive, HER2 negative |
| Luminal B | ER and/or PR positive, HER2 positive |
| Her2neu | ER and PR negative, HER2 positive |
| Basal like (Triple negative) | ER, PR and HER2 negative |

## Results

### Demographics and general characteristics of neoplasm

Among the patients who have undergone an excisional procedure for breast cancer, those belonging to the 46-60 years age bracket were the commonest, followed by the 31-45 years age group (24.29%). Only 08 patients (4.5%) belonged to the under 30 age group. Mean age of the cases was 51.39 years. MRM was the most frequent specimen received (73.60%), followed by excisions less than total mastectomy (14.04%). Overall, MRM was most frequently performed in the age group over 60 (86%), followed by 31-45 years (76%). Majority of surgeries performed in the under 30 age group was excisions less than total mastectomy (75%).

### Histological types of neoplasms

The commonest histological variant of breast carcinoma was found to be Infiltrating Ductal Carcinoma (Table 2), Not Otherwise Specified (IDCNOS) (83.62%); a few cases of mucinous carcinoma, papillary carcinoma, invasive lobular carcinoma (ILC) and medullary carcinoma were reported. Two cases of combined ductal and lobular carcinoma (IDC, ILC) was reported in this period. A rare case of neuroendcrine carcinoma breast was also identified, which turned out to be positive for ER as well as neuroendocrine markers Synaptophysin and Chromogranin. One metastatic oral squamous carcinoma in the breast was also reported, which was, however, excluded from the anaylsis. 09 (nine) cases were reported as having only Ductal Carcinoma in Situ (DCIS), without an invasive component. However, grading and immunohistochemistry have been performed on them.

**Table 2:** Histological types of breast carcinomas

| Type of carcinoma | Number |
| --- | --- |
| IDCNOS | 148 ( 83.62%) |
| DCIS | 9 ( 5.08%) |
| Mucinous | 5 ( 2.82%) |
| Papillary | 4 ( 2.26%) |
| Medullary | 3 ( 1.69%) |
| ILC | 3 ( 1.69%) |
| IDC, ILC | 2 ( 1.13%) |
| Metaplastic | 1 ( 0.56%) |
| Cribiform | 1 ( 0.56%) |
| Neuroendocrine | 1 ( 0.56%) |
| **Total** | **177 ( 100.00% )** |

### Histological grading

SRB score could be calculated in 161 cases, which were then placed in three grades (Figure 1). Majority of cancers were Grade 3 (47.8%), followed by Grade 2 (36.02%) and Grade 1 (16.15%). Grade 3 neoplasms were commonest in the age group above 60 (69%), followed by the 31-45 age group (55%). There was no statistically significant difference of grades of carcinoma in between age groups (χ^2^ = 13.67, p = 0.09)

**Figure 1:**
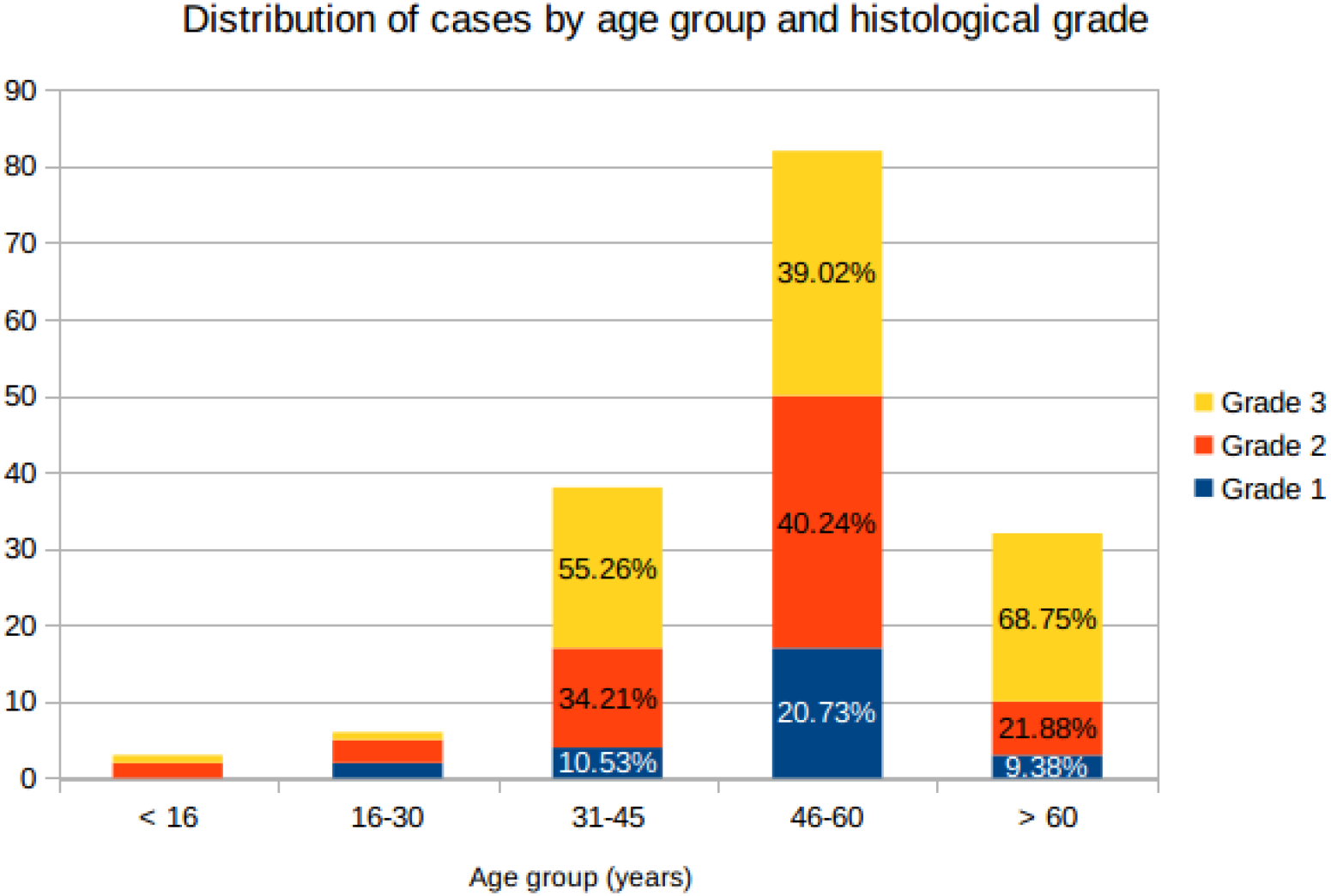
Age distribution

A one way ANOVA between the size of primary neoplasms (i.e. excluding those post NACT) between three grades (Figure 2) showed no significant association (F = 0.59, p = 0.55).

**Figure 2:**
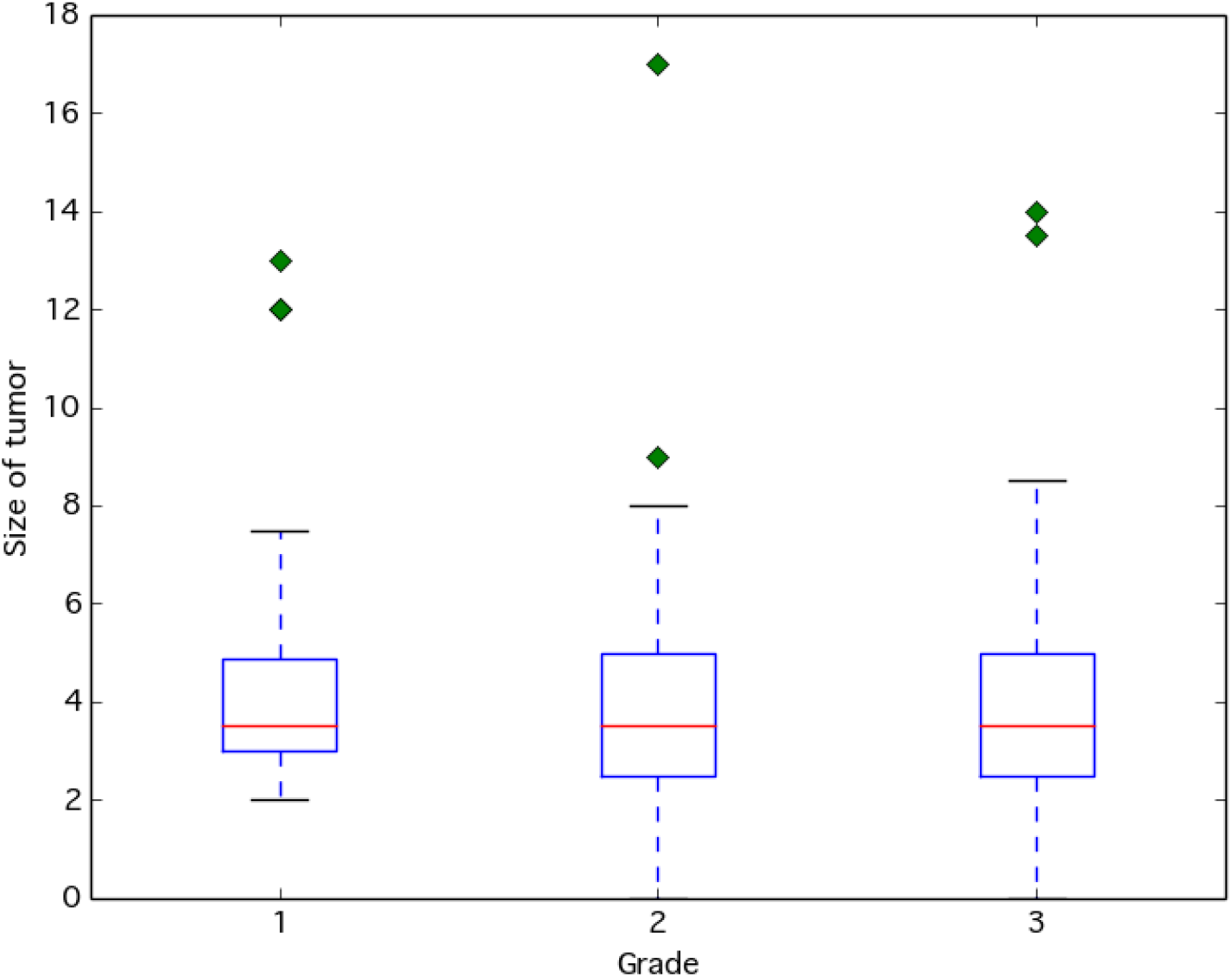
Distribution of tumor size by grade

### Stage at operation

T staging was performed on all cases except 01 block and slide received from another laboratory. Majority of cases were T2 (52.27%) at operation, followed by T3 (21.02%) and T1 (9.66%). 14 cases (7.95%) were in stage T4 at operation. Among cases not exposed to NACT, grade 3 neoplasms were proportionately more frequent to present at stage T3 (53.85%). There was no significant association between grade and T stage of neoplasms (exlcuding cases with prior NACT) (χ^2^ 9.58; p=0.29), as seen in Figure 3.

**Figure 3:**
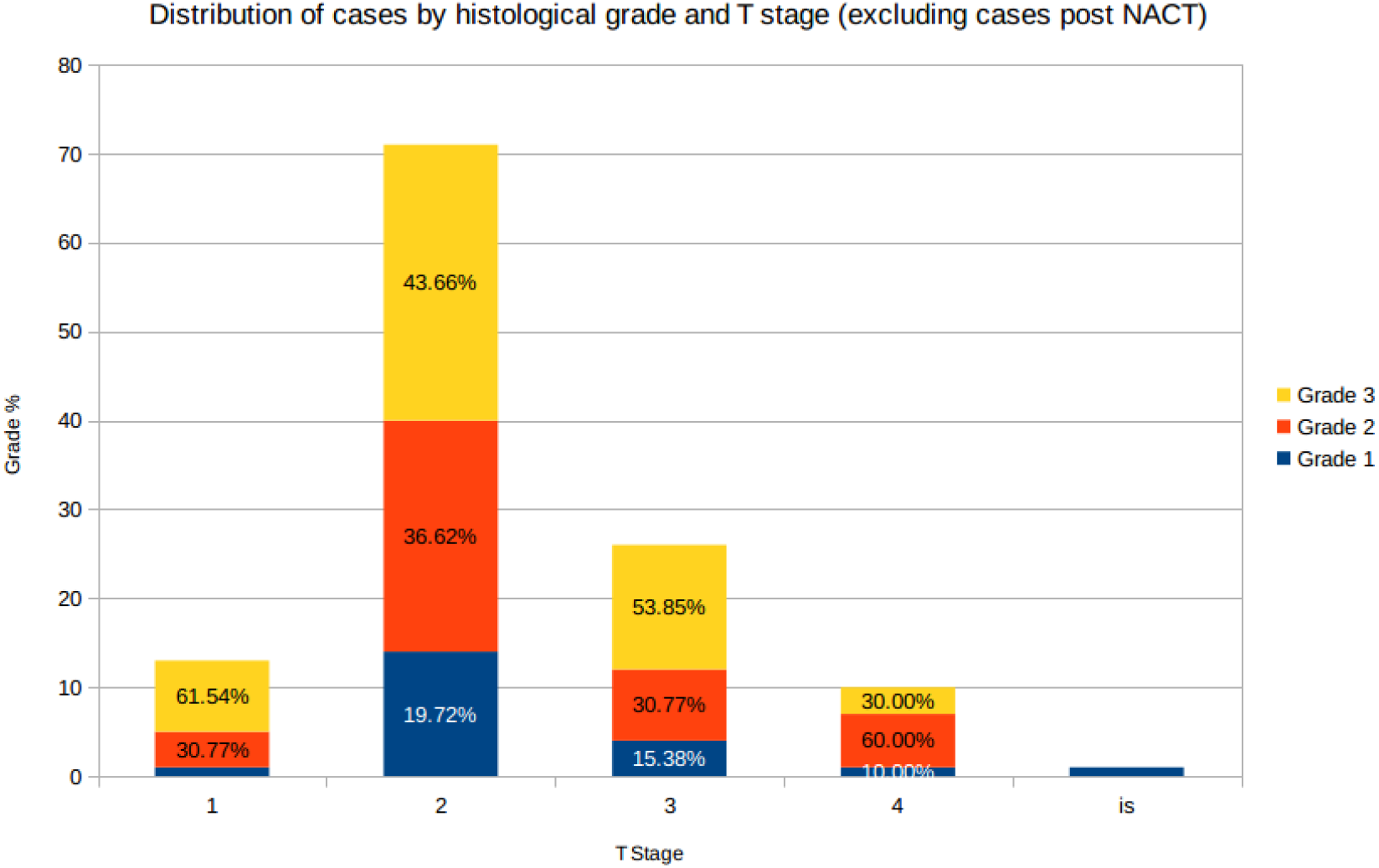
Distribution of grade by T stage

Higher stage at presentation (T3 and T4) was commonest in the age groups 31-45 and 46-60 years (exlcuding cases treated with prior NACT, N = 136) (Figure 4). T stage at presentation (excluding Tx) showed no significant difference between age groups. (χ^2^ 18.81, p = 0.09) (excluding Tx). T staging was not done in blocks received from other labs.

**Figure 4:**
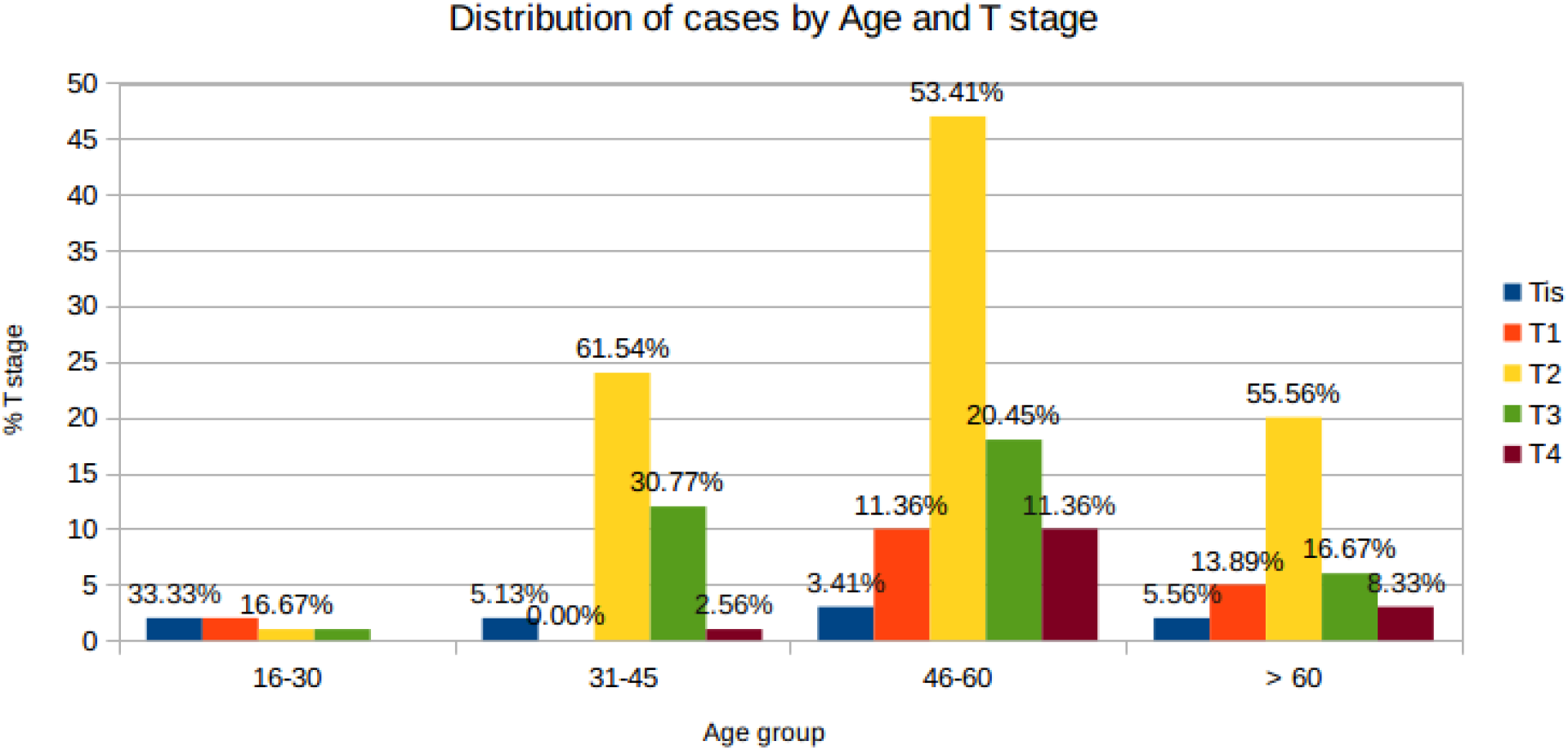
Distributiio by age aod T stage

### Lymph nodes

On an average, 14.16 lymph nodes were dissected from MRM with axillary dissections. Average number of lymph nodes (LN) involved was 3.34 lymph nodes. The mean percentage of lymph nodes involved 23.52 %, and of all MRM cases, 80 showed lymph node involvement (61.5%).

Table 3 shows percentage lymph node involvement (where >= 10 lymph nodes have been dissected) by grade of neoplasm. Majority of cases showing > 60% lymph node involvement (61.54%) belonged to grade 3 tumors. There was no significant difference of lymph node involvement between different grades of neoplasm (χ^2^ 0.99, p = 0.90).

**Table 3:** Distribution of cases by grade and percentage lymph node (LN) involvement (where > 10 lymph nodes were dissected)

| Percentage LN Involved | Grade 1 | Grade 2 | Grade 3 | Total |
| --- | --- | --- | --- | --- |
| < 30% | 10 (12.20%) | 29 (35.37%) | 43 (52.44%) | 82 (100%) |
| 30-60% | 2 (9.52%) | 8 (38.10%) | 11 (52.38%) | 21 (100%) |
| > 60% | 2 (15.38%) | 3 (23.08%) | 8 (61.54%) | 13 (100%) |
| **Total** | 14 ( 12.07% ) | 40 ( 34.48% ) | 62 ( 53.45% ) | 116 (100%) |

There is a significant association between lymphovascular invasion (LVI) and N stage (χ^2^ = 15.49, p = 0.001), as seen in Table 4. However, LVI is not statistically associated with grade of neoplasm (χ^2^ 2.30, p = 0.31).

**Table 4:** Distribution of cases by presence of lymphovascular invasion (LVI) and N stage

| N Stage | LVI Absent | LVI Present | Total |
| --- | --- | --- | --- |
| 0 | 73 (90.12%) | 8 (9.88%) | 81 (100%) |
| 1 | 36 (75.00%) | 12 (25.00%) | 48 (100%) |
| 2 | 22 (66.67%) | 11 (33.33%) | 33 (100%) |
| 3 | 8 (53.33%) | 7 (46.67%) | 15 (100%) |
| **Total** | 139 ( 78.53% ) | 38 ( 21.47% ) | 177 (100%) |

### Immunohistochemical parameters

In the cases where immunohistochemistry (IHC) was performed (N=108), stratification was done in the following manner (Table 5).

**Table 5:** Distribution of cases by immunohistochemical categories (where immunohistochemistry was performed)

| Luminal B | Her2 only | Triple negative | Total |
| --- | --- | --- | --- |
| 6 (5.56%) | 16 (14.81%) | 41 (37.96%) | 108 (100%) |

There was no significant difference in age group (p = 0.18), histopathologic type (p = 0.54), T stage (p = 0.10), N stage (p = 0.70), histopathologic grade (p = 0.44), desmoplastic reaction (p = 0.67), skin involvement (p = 0.73), mitotic count (p = 0.68), lymphovascular invasion (p = 0.36) and percentage of lymph node involvement (p = 0.32) between immunohistochemical categories. As individual markers, ER, PR and HER2 expression was not significantly associated with grade (p = 0.07, 0.87 and 0.69 respectively). Table 6 shows the distribution of cases by IHC categories and morphological criteria. Figure 6 shows a core biopsy of a neoplasm which was classified as ER negative, PR positive and HER2 negative.

**Table 6:** Distribution of cases by immunohistochemical categories and morphological parameters (where IHC was performed)

| T Stage (excluding Tx) | Luminal B | Luminal A | Her2 only | Triple negative | Total |
| --- | --- | --- | --- | --- | --- |
| 1 | 1 (8.33%) | 8 (66.67%) | 0 | 3 (25.00%) | 12 (100%) |
| 2 | 2 (3.33%) | 26 (43.33%) | 9 (15.00%) | 23 (38.33%) | 60 (100%) |
| 3 | 3 (14.29%) | 5 (23.81%) | 6 (28.57%) | 7 (33.33%) | 21 (100%) |
| 4 | 0 | 3 (37.50%) | 0 | 5 (62.50%) | 8 (100%) |
| is | 0 | 0 | 1 (25.00%) | 3 (75.00%) | 4 (100%) |
| **Total** | 6 ( 5.71% ) | 42 ( 40.00% ) | 16 ( 15.24% ) | 41 ( 39.05% ) | 105 (100%) |
| N Stage (where N stage could be assessed) |  |  |  |  |  |
| 0 | 1 (2.04%) | 23 (46.94%) | 7 (14.29%) | 18 (36.73%) | 49 (100%) |
| 1 | 2 (8.33%) | 9 (37.50%) | 2 (8.33%) | 11 (45.83%) | 24 (100%) |
| 2 | 2 (7.41%) | 9 (33.33%) | 5 (18.52%) | 11 (40.74%) | 27 (100%) |
| 3 | 1 (12.50%) | 4 (50.00%) | 2 (25.00%) | 1 (12.50%) | 8 (100%) |
| **Total** | 6 ( 5.56% ) | 45 ( 41.67% ) | 16 ( 14.81% ) | 41 ( 37.96% ) | 108 (100%) |
| Grade (where grading was possible) |  |  |  |  |  |
| Grade 1 | 1 (16.67%) | 3 (50.00%) | 1 (16.67%) | 1 (16.67%) | 6 (100%) |
| Grade 2 | 1 (3.57%) | 16 (57.14%) | 4 (14.29%) | 7 (25.00%) | 28 (100%) |
| Grade 3 | 4 (5.71%) | 26 (37.14%) | 10 (14.29%) | 30 (42.86%) | 70 (100%) |
| **Total** | 6 ( 5.77% ) | 45 ( 43.27% ) | 15 ( 14.42% ) | 38 ( 36.54% ) | 104 (100%) |

| LN Involved Percentage (where lymph node dissection was done) |  |  |  |  |  |
| --- | --- | --- | --- | --- | --- |
| < 30% | 3 (4.92%) | 27 (44.26%) | 9 (14.75%) | 22 (36.07%) | 61 (100%) |
| 30-60% | 1 (6.67%) | 2 (13.33%) | 2 (13.33%) | 10 (66.67%) | 15 (100%) |
| > 60% | 2 (13.33%) | 8 (53.33%) | 3 (20.00%) | 2 (13.33%) | 15 (100%) |
| **Total** | 6 ( 6.59% ) | 37 ( 40.66% ) | 14 ( 15.38% ) | 34 ( 37.36% ) | 91 (100%) |
| Lymphovascular invasion |  |  |  |  |  |
| Absent | 3 (4.00%) | 35 (46.67%) | 10 (13.33%) | 27 (36.00%) | 75 (100%) |
| Present | 3 (9.09%) | 10 (30.30%) | 6 (18.18%) | 14 (42.42%) | 33 (100%) |
| **Total** | 6 ( 5.56% ) | 45 ( 41.67% ) | 16 ( 14.81% ) | 41 ( 37.96% ) | 108 (100%) |
| Mitotic score (where SRB score could be calculated) |  |  |  |  |  |
| 1 | 1 (7.69%) | 7 (53.85%) | 2 (15.38%) | 3 (23.08%) | 13 (100%) |
| 2 | 2 (5.13%) | 20 (51.28%) | 4 (10.26%) | 13 (33.33%) | 39 (100%) |
| 3 | 3 (5.77%) | 18 (34.62%) | 9 (17.31%) | 22 (42.31%) | 52 (100%) |
| **Total** | 6 ( 5.77% ) | 45 ( 43.27% ) | 15 ( 14.42% ) | 38 ( 36.54% ) | 104 (100%) |

### Hormone receptors in post NACT and primary tumors

Table 7 shows that there is a significant difference in ER, PR expression between cases post NACT and primary cases (χ^2^ = 5.66; p = 0.017); in particular, NACT reduces hormone receptor expression. However, there was no such difference in Her2 expression (χ^2^ = 2.28, p =0.13).

**Table 7:**
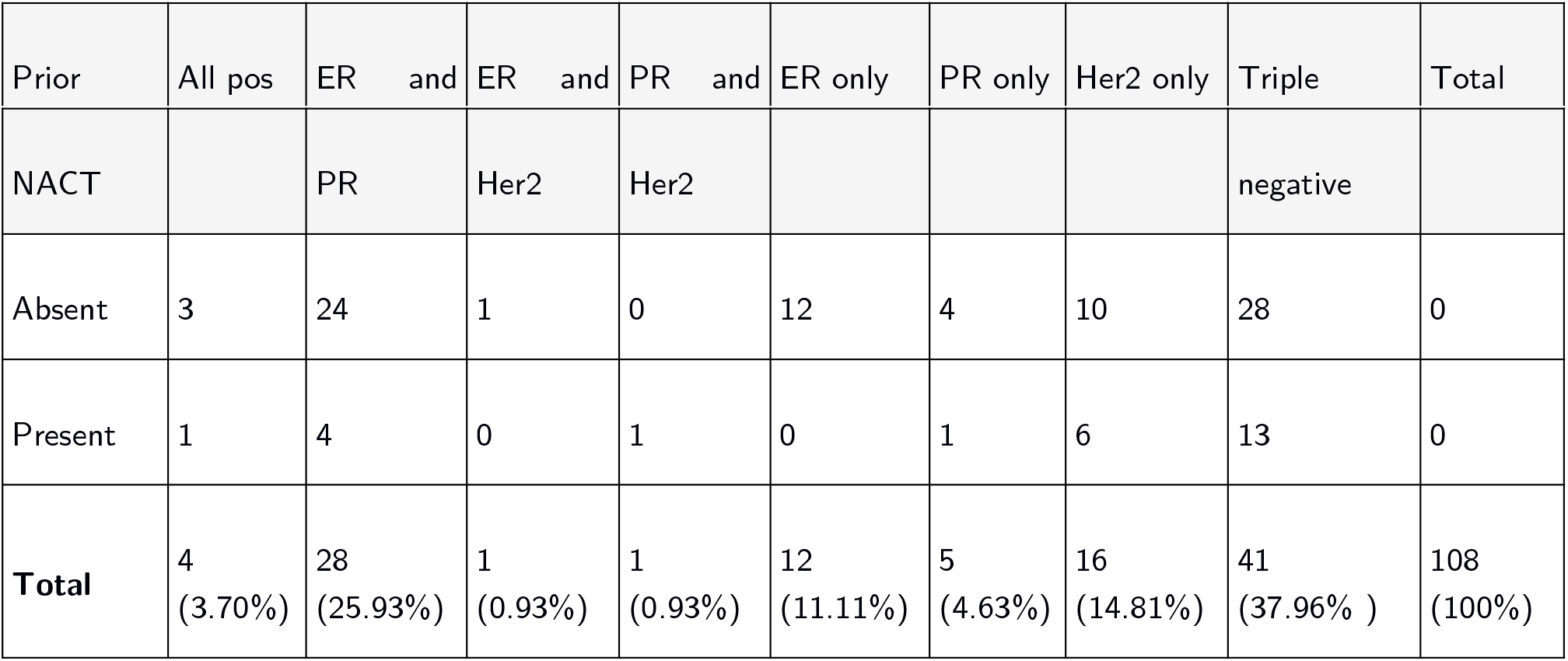
Distribution of cases by prior NACT and IHC markers

Of the five cases which have recurred during the study period, only 01 is triple negative for ER, PR and HER2. 01 case was positive for HER2, and the rest of three cases were positive for ER and/ or PR. However, on recurrence, one other case has lost hormone receptor expression and become triple negative.

### Pre and Post NACT IHC expression

A number of studies have shows that neo adjuvant chemotherapy (NACT) significantly changes the ER, PR and Her2 expression in breast cancers, [8]; usually, marker positive neoplasms turn marker negative [9]. In the present study, 19 (nineteen) patients were followed up) from core biopsy to mastectomy after NACT. Figure 5 shows the effects of NACT on expression of hormone receptors: average reduction in HER2 score was 0.21, reduction in ER score 1.57, and reduction in PR score 0.57 (Figure 5).

**Figure 5:**
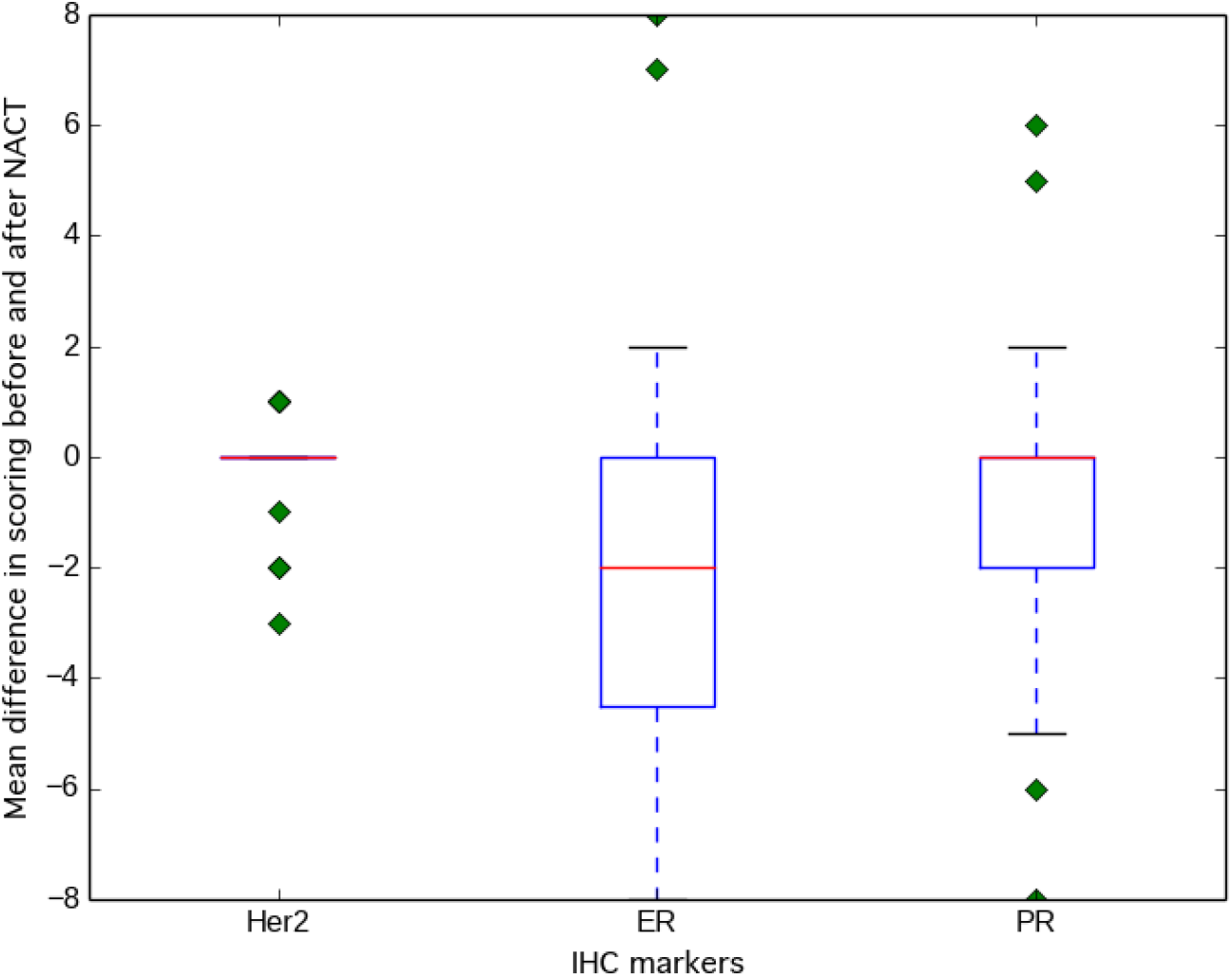
Difference in IHC markers post NACT

## Discussion

Breast cancer is a heterogeneous disease; attempts to predict its molecular nature and clinical behavior through phenotypic markers such as immunohistochemistry has been made. In the present study, we have looked at morphological parameters which determine the clinical behavior of a neoplasm, and their correlation, if any, with immunohistochemical subtypes.

The mean age of patients in this study was 51.39 years, which is similar to previous studies in India [10] [11]. In one of the studies in India from 2013 of 50 patients [12] it was found that majority of patients had operable breast cancer at presentation. Most of the patients were between 40-60 yr of age which suggest shift towards younger age groups. Hormone receptor positivity was seen in two third of the patients. Small size, low grade and tumours without axillary lymph node metastasis showed more frequent receptor positivity. According the Consolidated Report of the Hospital Based Cancer Registries 2012 - 2014, 48% of all breast cancer patient in India present below 50 years of age [13]. In the present study 73 patients (41.01%) presented below the age of 50 years. 4.49% of patients presented at an age less than 30.

In a large study from Delhi in 2005 [11], the Infiltrating duct carcinoma (IDC) was found to be commonest type (88.2%), followed by infiltrating lobular carcinoma (ILC) (3.7%). In the present study majority of cases were IDCNOS, most of which presented in Stage T2 (54%). Majority of the two other variants (mucinous, papillary) also presented in stage T2. The findings are similar to a large case series of 135157 cases published in 2005 [14], which reported that mucinous carcinomas were less likely to present at an advanced stage. The same study also reported that mucinous carcinomas are more likely to be positive for ER/ PR. Among two mucinous carcinomas in this study in which IHC was performed, one was positive for all markers, another was triple negative. Notably, a case of neuroendocrine carcinoma was positive for ER as well as Synaptophysin and Chromogranin.

Majority of neoplasms in the present study were in Grade 3 (47.83%). Similar to the study by Kumar et al [10], majority of Grade 3 neoplasms were triple negative; only 01 triple negative neoplasm was grade 1. Unlike that study, no statistically significant association was found between grade of neoplasm and immunophenotypic category. The grade of the neoplasm was also unrelated to the size at presentation, age and T stage.

In the study by Saxena et al [11], T stage 3 (35.2%) was the commonest. However, in the series by Kumar et al [10], T2 was the most frequent (55.4%). The findings in the present study was similar to the later, T2 being the commonest stage at presentation (54.44%). 12 out of 41 triple negative neoplasms (29.26%) were in stage T3 or higher; triple negative neoplasms were also the commonest in T stage 3 and 4. The finding is similar to Kumar et al [10], who have found 94.1% of triple negative neoplasms in stage T2 or higher; the corresponding figure in the present study being 92.68%. However, there was no statistically significant difference in T stage between different immunohistochemical categories.

In the study by Saxena et al, lymph node positivity was observed in 80.2% of cases; in the present study it was significantly lower (61.5%). No statistically significant association was noted between N stage and immunohistochemical category. The highest proportion of lymph node involvement was seen in the Luminal B subtype (50% cases involving > 60% of dissected lymph nodes); among the cases with < 30% lymph node involvement, Luminal A subtype was commonest (43.64%), which is similar to the study by Kumar et al [10]. However, there was no statistically significant association between lymph node involvement and immunohistochemical subtypes. The highest incidence of LVI was seen in Luminal B subtype (75%), which is similar to the study by Kumar et al [10]. There was no significant difference in LVI between immunohistochemical categories.

In the division of neoplasms into immunohistochemical categories, we have followed the classification proposed by the Carolina Breast Cancer Study [6]. In a study in 2015 of 56 patients [10], the Luminal A subtype (ER and/ or PR positive, HER2 negative) was most prevalent (34%), followed by Triple negative subtype (25%). Luminal B (ER and/ or PR positive, HER2 positive) and HER2 subtypes (isolated HER2 positive) had same prevalence i.e. 18%. Histological grade and ER negative status showed strong correlations with basal markers. In a smaller study of 50 breast cancers the authors concluded that hormone positive tumors are generally of good prognosis. [15]. In the present study, Luminal A subtype was the most frequent (41.67%), followed by the triple negative subtype (37.96%). This is in keeping with the study by Kumar et al [10]. However, unlike previous studies in India, there was no statistically significant difference in morphologic criteria (type of neoplasm, grade, size, dense infammation, desmoplasia, lymphovascular invasion, nodal involvement, mitotic count) between different immunohistochemical categories.

Among the neoplasms which were received post NACT, hormone receptor (ER/ PR) expression was significantly reduced, but not HER2.Only 01 case, among the five which had recurred, was triple negative.

Among the cases followed up during the period of study, maximal reduction was seen in ER expression after NACT, followed by PR expression. There was only minimal loss of HER2 expression. This is in consonance with the study by Ozmen et al [16], who reported a reduction in ER/PR expression in 28% cases, but HER2 reduction only 3% cases.

## Conclusion

We conclude that immunohistochemical categories do not show a statistically significant association with morphologic parameters of breast neoplasms (size, grade, stage, mitotic count, desmoplasia, dense infiltrate, lymphovascular invasion). However, we have found association between lymphovascular invasion within the primary tumor and lymph node involvement. Our findings suggest that there might be additional genetic subtypes underlying the immunophenotypic variants which are hitherto unknown, but play a vital role in disease behavior. We feel the need to validate our findings over a prospective study with a larger sample size.

